# Choice of lipid supplementation for *in vitro* erythroid cell culture impacts reticulocyte yield and characteristics

**DOI:** 10.1101/2025.07.11.664344

**Authors:** CM Freire, NR King, M Dzieciatkowska, D Stephenson, J.G.G Dobbe, GJ Streekstra, A D’Alessandro, TJ Satchwell, AM Toye

**Affiliations:** School of Biochemistry, University of Bristol, Bristol, UK; Department of Biochemistry and Molecular Genetics, University of Colorado Anschutz Medical Campus, Aurora, CO 80045, USA; Amsterdam UMC location University of Amsterdam, Biomedical Engineering and Physics, Meibergdreef 9, Amsterdam, the Netherlands; Centre for Biomedical Research, School of Applied Sciences, University of the West of England, Bristol, UK

## Abstract

Lipids, particularly cholesterol, are critical components of red blood cell (RBC) membranes, influencing protein function, cell stability, and deformability. Reticulocytes (young RBC) derived from *in vitro* erythroid cultures have been reported to possess less cholesterol than their native counterparts, compromising their functional integrity and lifespan. However, variability in starting materials and culture protocols between studies has hindered definitive conclusions regarding the nature and consequences of this lipid deficiency.

Here, we evaluated the influence of lipid sources on reticulocyte quality using a well-established CD34⁺ erythroid culture system. We compared the use of human AB-serum and Octaplas (solvent/detergent-treated pooled plasma) as lipid sources. Our results reveal that detergent-treated plasma leads to cholesterol-deficient reticulocytes with impaired characteristics, including reduced filtration yield, heightened osmotic fragility, and altered PIEZO1 activity. In contrast, AB-serum supported the generation of functionally stable reticulocytes, with cholesterol supplementation required to rescue the defects observed with plasma.

Importantly, this study provides the first integrated lipidomic, metabolomic, and proteomic characterisation of *in vitro*-derived reticulocytes cultured under distinct lipid conditions. These multi-omic datasets offer new insights into the consequences of reduced lipid availability during erythroid culture and offer new insights into how culture media affects the development and functionality of lab grown blood.

## Introduction

Red blood cells (RBCs) are continuously subjected to shear stress during circulation and have to deform repeatedly to traverse narrow capillaries and splenic slits^1^. This exceptional deformability is largely due to their highly specialised membrane, which comprises a spectrin based cytoskeleton anchored to a cholesterol rich lipid bilayer. Unlike most cellular membranes which typically contain 10–30% cholesterol, the RBC membranes contain up to 50% cholesterol out of total membrane lipids^2^, a feature critical for maintaining mechanical stability, fluidity and overall function.

The connection between cholesterol and erythrocyte physiology has been recognised for over a century since cholesterol was first identified in red cell membranes. Studies have since shown cholesterol levels influence RBC deformability^3,4^ and are linked to haematological disorders associated with altered blood ^5^viscosity. Dysregulation of cholesterol homeostasis affects RBC lifespan, clearance, and oxygen transport efficiency^6–8^. For example, foetal erythropoiesis involves up-regulation of cholesterol metabolism ^9^, and is linked to activation of p53-dependent activation of cholesterol uptake by ABCA1 and suppression of cholesterol synthesis via the mevalonate pathway^10^. This regulatory axis is essential during adult erythropoiesis, where p53 influences iron uptake by controlling the transcription of the ferrireductase Steap3^11^. Hypercholesterolemia has also been reported as disrupting RBC functional and structural properties, by accumulating cholesterol within the plasma membrane cell deformability is reduced, impacting oxygen transport^12^. In mature RBCs, cholesterol depletion, which occurs during refrigerated storage, has been linked to changes in morphology, increased membrane rigidity^13^, reduced deformability and, enhanced splenic clearance^14^. These findings highlight the importance of maintaining cholesterol availability during *in vitro* erythroid culture, where lab-grown reticulocytes must emulate native RBCs membrane properties to survive and function effectively *in vivo*.

The heightened interest in lab-grown RBCs as potential blood substitutes has intensified the focus on the characteristics of cultured reticulocytes and how closely they mimic their native counterparts. Previous studies have shown that lab-grown reticulocytes exhibit similar blood group antigen expression to their originating donor cells^15,16^, possess a proteomic profile comparable to native reticulocytes^17,18^, and are capable of deforming^19,20^. Furthermore, genetic modifications introduced into cultured erythroid cells to mimic patient specific mutations yield reticulocytes that display the characteristic phenotype expected, provided the phenotype manifests at the immature RBC stage^21–23^. These findings suggest that lab-grown reticulocytes possess requisite membrane protein structures and interactions, supporting their potential as functional RBC substitutes.

One currently underexplored factor influencing the properties of lab grown blood is their plasma membrane lipid composition. Outside the highly regulated and nurturing *in vivo* bone marrow microenvironment, the impact of exogenously supplied lipid sources used during erythroid culture is of particular importance. In the RESTORE clinical trial, which is currently evaluating the performance of lab-grown reticulocytes compared to donor derived red blood cells in healthy volunteers, AB-serum (alongside human serum albumin (HSA)) is used as a lipid source and no detrimental effects observed on the characteristics of the lab-grown reticulocytes. In contrast, several studies ^20,24,25^ have reported cholesterol deficiencies in cultured reticulocytes, which required supplementation to preserve reticulocyte integrity. These discrepancies are often difficult to interpret due to inherent variability in experimental reagents, donor cell sources and culture protocols. Nevertheless, we hypothesised that the observed differences may be attributable to the lipid sources employed-particularly as the latter studies all used detergent/solvent extracted plasma. Addressing this question requires a more standardised and systematic comparison of lipid supplements used in erythroid culture systems.

In order to systematically assess the lipid content of cultured reticulocytes produced using distinct lipid sources, lipids were provided through supplementation of *in vitro* erythroid cultures with AB-serum, or detergent/solvent-extracted (S/D) plasma in the presence or absence of cholesterol supplementation. Our findings demonstrate that the choice of lipid sources in erythroid culture significantly affects cholesterol availability, which in turn affects cell yield and reticulocyte characteristics. Moreover, we provide the first comprehensive lipidomic, metabolomic, and proteomic comparisons of reticulocytes derived from these diverse lipid sources.

## Results

### Reticulocyte yield is impacted by choice of lipid source in differentiation media

To elucidate the role of lipid supplementation in *in vitro* erythroid cultures, multiple lipid sources were tested in adult hematopoietic stem cell (HSC) cultures. We used a well-established three-phase culture method comprised of Iscove’s Modified Dulbecco’s Medium (IMDM) supplemented with erythropoietin (EPO), insulin, heparin, holo-transferrin, as base media^31^. The culture media composition was kept identical but the lipid sources used were either 3% AB-serum and 2% HSA (called here AB-serum) or 5% human pooled plasma (brand name Octaplas®, referred to as Plasma for simplicity), or plasma supplemented with 50mg/L cholesterol-rich-lipids (Plasma + CRL). **Figure 1A** summarises the culture conditions tested, and the culture protocol followed. Cells were cultured for 20 days with **Figure 1B** illustrating the cumulative fold expansion of the erythroid cultures in the three tested media compositions. The standard deviation between the 6 donors tested is high, particularly when cultured in AB-serum, most likely due to donor biological variability in terms of progenitor expansion. The AB-serum cultured cells tend to experience a higher proliferation from day 5 onwards compared to the plasma culture, plateauing at 4.8±4.5x10^4^-fold at day 15. Plasma-cultured cells on average exhibited only 1.9±1.1 x10^4^-fold proliferation on day 15 and the supplementation with CRL increased proliferation to 3.6±2.4 x10^4^-fold on day 18.

**Figure 1:**
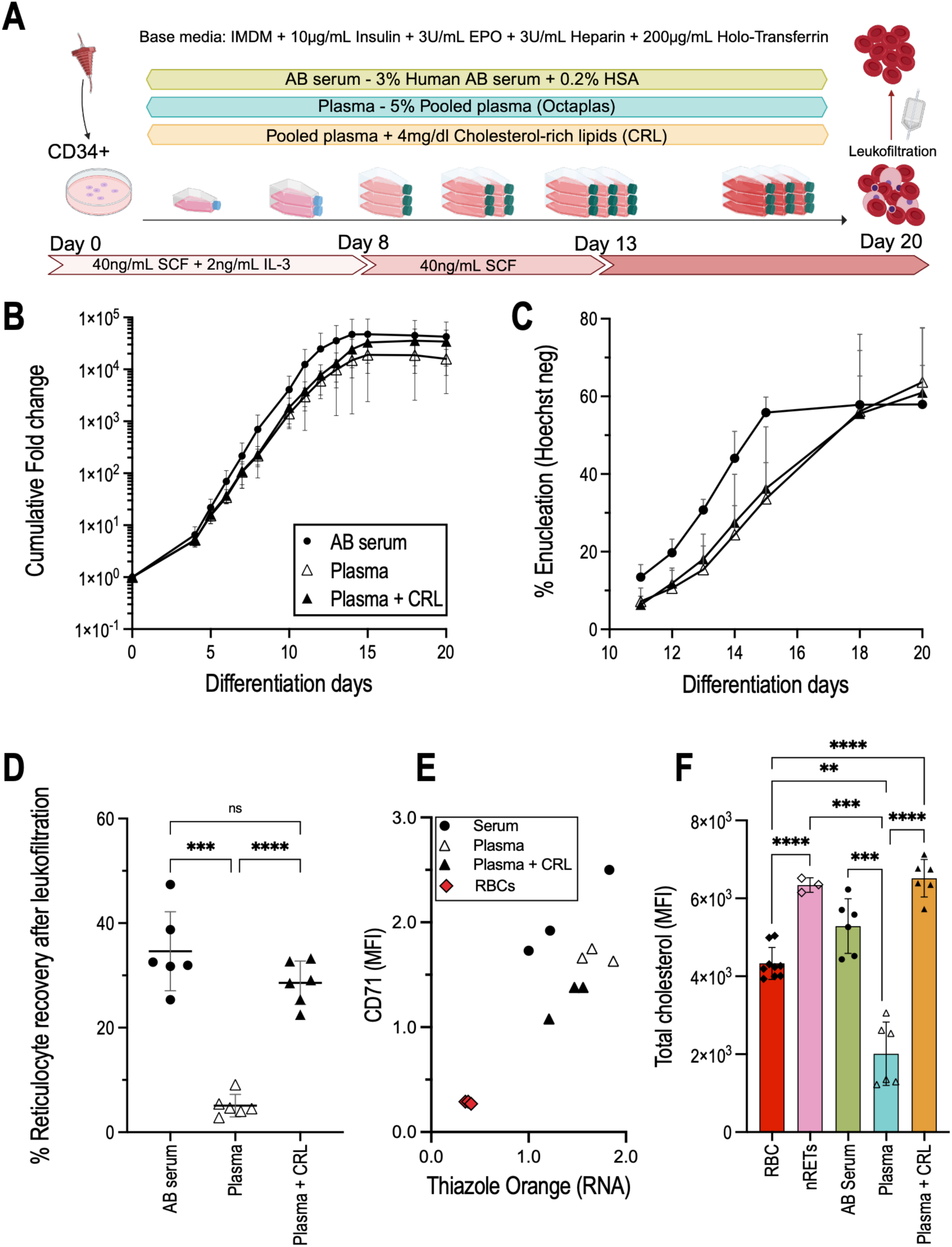
Lipid source used in primary *in vitro* erythropoiesis affects cell expansion, enucleation rate, and filtration yields. **A** CD34-positive cells (human hematopoietic stem cell marker) are isolated from leukocyte reduction filters and cultured for 20 days using a three-step culture protocol. The base media consists of Iscove’s Modified Dulbecco’s Medium (IMDM) supplemented with insulin, erythropoietin (EPO), heparin, holo-transferrin, and either AB serum or Plasma as lipid sources. The addition of cholesterol-rich lipids (CRL) to plasma cultures was also explored. In the first stage, days 0-7, media is supplemented with stem-cell factor (SCF) and interleukin-3 (IL-3). The second stage until day 13 depends only on SCF supplementation and the third being only base media. On day 20 cells are leukofiltered to obtain a pure reticulocyte population. **B** Cumulative fold expansion (log_10_ scale) of adult CD34+ cells differentiation grown in media containing either human AB serum, pooled human plasma, or pooled plasma supplemented with Cholesterol-Rich-lipids (CRL). **C** Percentage enucleation from day 11 of differentiation. Reticulocytes were identified by Hoechst negativity. n=6, error bars represent the standard error of the mean. **D** Percentage of reticulocytes recovered after leukofiltration on day 20 of erythroid differentiation. A parametric (normality confirmed with a Shapiro-Wilk test) Brown-Forsythe and Welch test was performed to test for differences between groups. p < 0.05 was considered statistically significant. n=6, error bars represent the standard deviation. **E** Median fluorescent intensity of CD71 (transferrin receptor) plotted against Thiazole Orange (RNA content) of filtered reticulocytes. Individual values for each donor (n=3) are represented, as well as three control red blood cell samples. Shaded regions correspond to a 95% confidence level calculated using a linear regression model in Rstudio. F Total cholesterol of matched samples (to panel A) was measured by flow cytometry using a filipin stain to associate osmotic fragility with cholesterol levels. The values were normalised to the RBCs. A parametric Brown-Forsythe and Welch test was performed to test for significance.

From days 11 to 20, enucleation was assessed on the basis of Hoechst negativity (**Figure 1C**). Until day 18 the percentage of reticulocytes in the AB-serum culture is greater than both Plasma cultures, indicating a faster progression through differentiation. Values remain constant for the AB-serum cultures day 15 onwards, contrary to Plasma which show a steady increase until day 18. Despite these differences, all three conditions yielded similar enucleation values at the time of harvest of the culture, an average of 61.0% ±2.9% (standard deviation, SD). Notably, supplementation with cholesterol does not alter enucleation percentages. For the same culture media composition containing AB-serum, values between 10^4^ and >10^5^-fold expansion and >60% enucleation have been reported^26,31^, consistent with the data collected here.

At the end of differentiation, the culture consists of a mixture of reticulocytes, pyrenocytes, and nucleated cells (mostly orthochromatic erythroblasts). To obtain a pure sample of reticulocytes (>98%) leukocyte reduction filters can be used to isolate the reticulocytes. Leukofilters are normally used after whole blood donations to eliminate white blood cells prior to transfusion, relying on the deformability of RBCs to easily pass through the filter while the stiffer nucleated cells are retained^32^. The cultures were filtered on day 20 and the yield (**Figure 1D**) was calculated based on total pre- and post-filtration cell counts. After filtration, the AB-serum culture condition averaged a yield of 34.6 ±7.5% (SD) while Plasma-cultured cells only achieved 5.1 ±2.2% reticulocyte recovery, a significant difference highlighting the decreased deformability of reticulocytes grown in Plasma. The addition of cholesterol-rich lipids to the plasma containing media rescued filterability levels to 28.9 ±4.1%.

The degree of reticulocyte maturation of the filtered cultures was next assessed through flow cytometry (**Figure 1E**) by plotting the expression of the transferrin receptor (CD71), a marker absent on mature erythrocytes, against Thiazole Orange – measuring residual RNA^33^. The individual data points indicate a higher CD71 expression, as judged by comparing MFI on AB-serum derived reticulocytes (RET^AB-serum^), compared with Plasma-derived (RET^Plasma^), either with or without cholesterol supplementation (RET^Plasma+CRL^).

Next, total cholesterol was assessed by Filipin stain (**Figure 1F**). This indicated a significant increase in total cholesterol of native RETs (nRETs) and RET^AB-serum^ compared with RBCs. RET^Plasma^ had a 54% decrease in total cholesterol when compared to RBCs (2.0±0.8 x10^3^ against 4.3±0.4 x10^3^), and 68% when compared to nRET (6.3±0.2 x10^3^). It should be noted that these results were obtained on the filtered reticulocytes, which only accounted for 5% of total reticulocytes present pre-filtration for plasma cultured cells. Upon cholesterol supplementation of plasma-cultured reticulocytes, total cholesterol levels increased to levels similar to that observed in nRETs. In fact, there is no significant difference between nRET, RET^AB-serum^, and RET^Plasma+CRL^, as all were higher than RBCs, although not significantly for RET^AB-serum^.

### Plasma grown erythroid cells have compensatory changes consistent with exposure to low cholesterol levels

The gene expression levels of key genes important for cholesterol homeostasis were evaluated using real-time quantitative PCR on RNA isolated from erythroblasts exposed to different lipid sources at day 8 of differentiation. Multiple genes known to control cholesterol biosynthesis were evaluated (**Figure 2A**) including the sterol-regulatory element-binding proteins 1 and 2 (*SREBP1* and *SREBP2*) expression (a transcription factor that controls cholesterol and lipid homeostasis), alongside 3-hydroxy-3-methylglutaryl-coenzyme A reductase (*HMGCR*) and Squalene monooxygenase (*SQLE*) which are both rate limiting enzymes in the cholesterol biosynthetic pathway. We also evaluated low-density lipoprotein receptor (*LDRL*) expression which plays a crucial role in cholesterol uptake.

**Figure 2:**
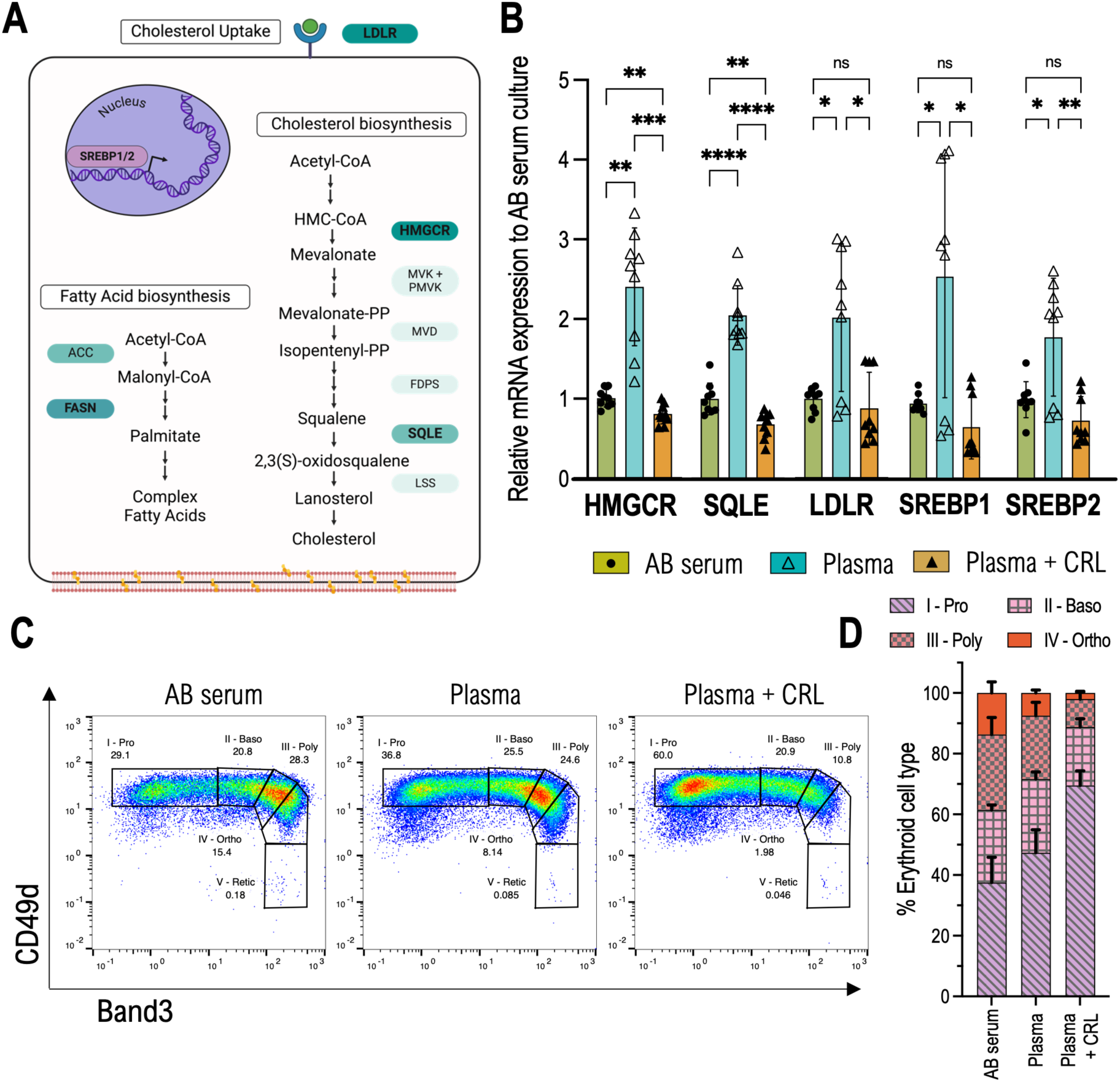
qPCR shows cholesterol biosynthesis is significantly increased in plasma cultured cells. **A** Main steps in cholesterol and fatty acid biosynthesis. Cholesterol biosynthesis begins with acetyl-CoA and proceeds via HMG-CoA reductase (HMGCR), forming mevalonate and downstream intermediates (mevalonate-PP, isopentenyl-PP, and squalene). Squalene epoxidase (SQLE) converts squalene to 2,3(S)-oxidosqualene, leading to lanosterol and cholesterol synthesis. Fatty acid biosynthesis also originates from acetyl-CoA, with acetyl-CoA carboxylase (ACC) producing malonyl-CoA, followed by fatty acid synthase (FASN) generating palmitate and complex fatty acids. SREBP1/2 transcription factors regulate these pathways, with SREBP1 driving fatty acid synthesis and SREBP2 controlling cholesterol biosynthesis. The LDL receptor (LDLR) mediates cholesterol uptake, maintaining lipid homeostasis. Made using Biorender. B mRNA levels of HMGCR (HMG-CoA reductase), SQLE (Squalene Epoxidase), LDLR (Low density lipoprotein receptor), SREBP1 and SREBP2 (Sterol regulatory element-binding protein 1 or 2) relative to GAPDH on day 8 of differentiation after CD34+ isolation. A parametric Brown-Forsythe and Welch test was performed to test for differences between groups. p < 0.05 was considered statistically significant (n=3, N=3). C Example waterfall plots of cells the same donor on day 8 of erythroid differentiation, labelled against Band3 and CD49d, grown in medium containing AB serum, Plasma, or Plasma supplemented with CRL. Corresponding percentage of each stage (I-Proerythoblast, II-Basophilic, III-Polychromatic, IV-Orthochromatic, and V-Reticulocyte) is indicated next to the corresponding gate. D Shows percentage of erythroid cell types present on day 8 for each condition studied, quantified as displayed in panel above.(n=3). Error bars represent the standard deviation.

Gene expression was significantly increased in the Plasma condition for all genes analysed (**Figure 2B**) and was either similar or reduced upon CRL supplementation when compared to AB-serum. *HMGCR*, *SQLE*, *LDLR*, and *SREBP1* all show greater than 2-fold increase in expression (2.4±0.74, 2.1±0.38, and 2.0 ±0.94, 2.7 ±1.6 respectively). *SREBP2* had a 1.8±0.76 increase in Plasma culture compared with AB-serum. However, it is notable one donor exhibited a less pronounced response, for all genes except SQLE, to the low cholesterol environment, suggesting that that there may be inherent variability among donors. When CRL supplementation occurs alongside plasma, all genes have a mean fold-expansion inferior to 1 in the Plasma+CRL condition, indicating lower gene expression than when cultured in AB-serum.

To control for possible differences in gene expression associated with differentiation progression the different cell stages of erythropoiesis were assessed by observing the expression of membrane proteins Band 3 and CD49d, with Band 3 increasing during terminal differentiation while CD49d decreases^34^. Using one donor as an example, **Figure 2C** exemplifies the gating strategy used to determine the percentages of erythroid cell type (Proerythroblast (I), Basophilic Erythroblast (II), Polychromatic Erythroblast (III), and Orthochromatic Erythroblast (IV), Reticulocyte (V)) for tested conditions. The combined results for 3 donors are visible in **Figure 2D**.

Interestingly, **Figure 2D** shows that both plasma-containing conditions differentiated slower than AB-serum, with the latter having only 37.5±8.31% of total cells left as Proerythroblasts while Plasma-only had 47.3±7.57%, and Plasma + CRL 69.5±4.82%. This observation suggests that the restoration of gene expression levels observed in the Plasma+CRL cells to those comparable with AB-serum is not due to similarities in erythroid cell populations but rather a response to cholesterol availability.

### Proteomics, metabolomics and lipidomic studies

To more extensively explore the effects of media composition on the reticulocyte produced from the different lipid sources, samples of filtered reticulocytes and RBCs were subjected to multiomics analysis.

The proteomic dataset was analysed initially by normalising the Max Label Free Quantification values against donor-matched RBCs (**Figure 3A-B**). Many of the proteins highlighted where abundance significantly changes across the three conditions were associated with processes related to reticulocyte maturation and red cell-specific functionality, including membrane trafficking, carbon dioxide transport, and hydration regulation. Although, notably, Fatty acid lipid synthase (FASN) was detected as upregulated in the plasma culture reticulocytes. To more precisely dissect the differences induced by the extracellular environment, we performed a second analysis normalising instead to AB serum-cultured reticulocytes, allowing for the quantification of 1021 proteins, up from 500 when comparing against RBCs, effectively doubling the proteomic depth. This aligns with findings from Gautier et al. (2018)^35^ who identified 654 proteins as being reticulocyte specific. The Enrichr software tool was used to detect functional relationships between the protein groups. Of note was the identification of three proteins significantly upregulated in Plasma only-grown reticulocytes, observed in **Figure 3C** – Farnesyl pyrophosphate synthase (FDPS), FASN, and Diphosphomevalonate decarboxylase (MVN), all known to be involved in lipid and cholesterol biosynthesis. All three proteins are regulated by the transcription factors SREBP (Reactome pathway knowledgebase 2022, p=0.00002441), found to also be upregulated in the differentiating plasma-grown erythroid cells by RT-qPCR (**Figure 2B**).

**Figure 3:**
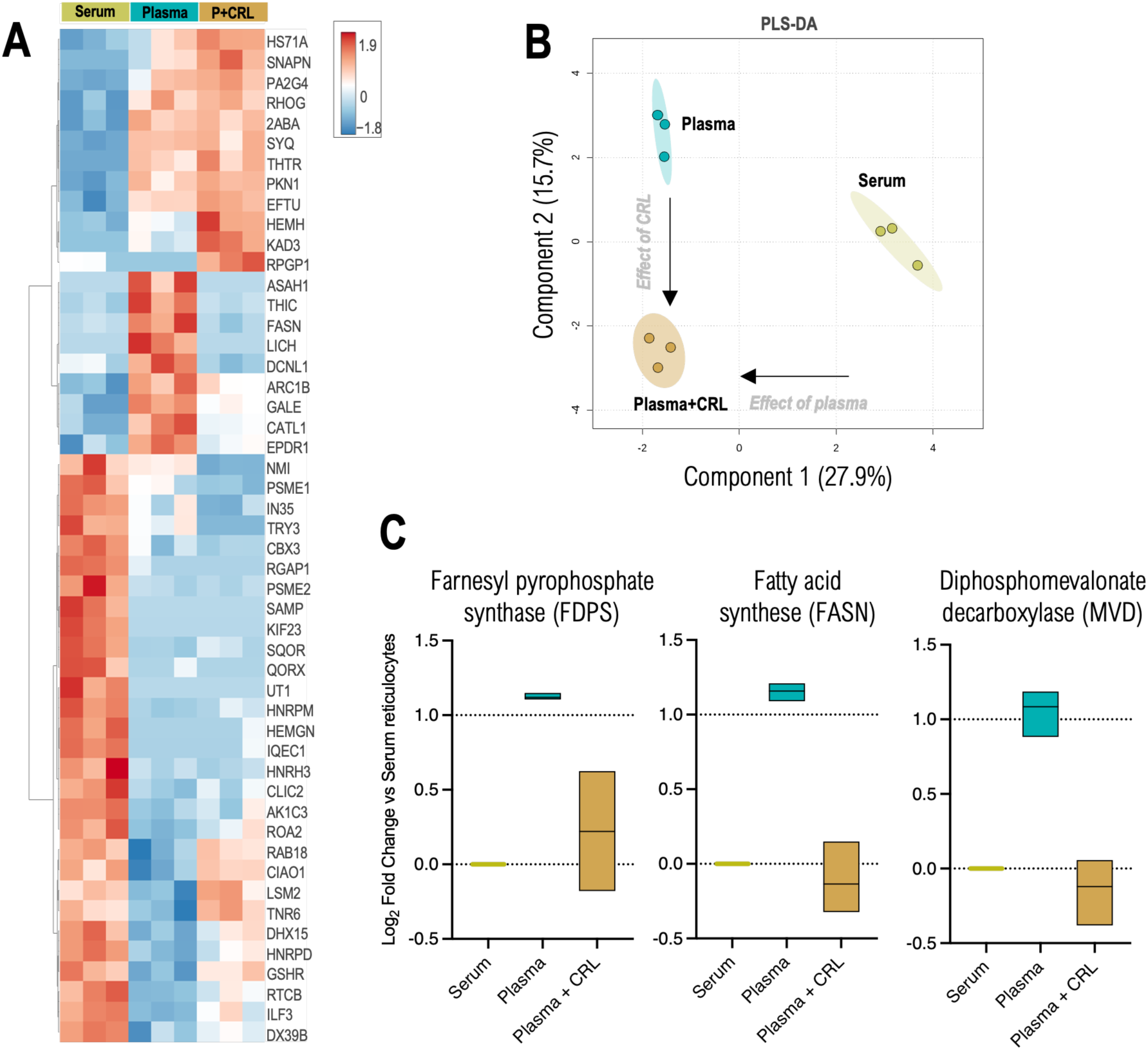
Summary of proteomic differences in CD34-derived reticulocytes grown in the presence of Serum, Plasma, or Plasma supplemented with cholesterol-rich lipids. Serum, Plasma and Plasma+CRL (P+CRL) compared with control RBCs (top 50 proteins by ANOVA in A and Partial least square-discriminant analysis – PLS-DA in B). C Box plots of upregulated proteins found in Plasma treated samples when compared with donor matched Serum. The log_2_(FC) was analysed for significance with non-parametric Kruskal-Wallis test with Dunn’s correction (p<0.05).

Metabolomics data was next analysed by comparing metabolites in Plasma or Plasma+CRL grown reticulocytes to AB serum reticulocytes. We observed that most metabolites (Supplemental Figure 1) have no significant alteration between culture systems and, even if observed to be significant these often translated into modest changes associated with a large degree of variability observed across the 3 donors analysed. Of note was the detected upregulation in UMP levels, particularly in the presence of cholesterol supplementation, and a reduction in pyridoxal. Decreases in succinate and citrate highlight alterations in the TCA cycle. The depletion of gamma-L-glutamyl-L-cysteine (precursor to glutathione^36^) and the increase in L-selenomethionine (common intermediate used for synthesising selenocysteine, essential for peroxidase activity^37^) also indicate alterations in antioxidant response.

A comprehensive lipidomics comparison was also conducted, and a summary heat map of the top 50 significant lipid changes identified between the cultured reticulocytes and matched RBCs is shown in **Figure 4A**. This identified several lipid classes with similar or higher abundance compared with red blood cells, which aligns with the fact that cultured reticulocytes must still lose around 20%^38^ of their membrane surface area during the maturation process. Acylcarnitines (AcCa, **Figure 4B**) are less abundant in serum-grown reticulocytes compared with both Plasma conditions, although this difference is not significant. Importantly, cholesterol levels were reduced in plasma, (Chol, **Figure 4C**) whereas cholesterol levels were observed to be significantly increased in both serum grown and Plasma supplemented with CRL reticulocytes relative to erythrocytes.

**Figure 4:**
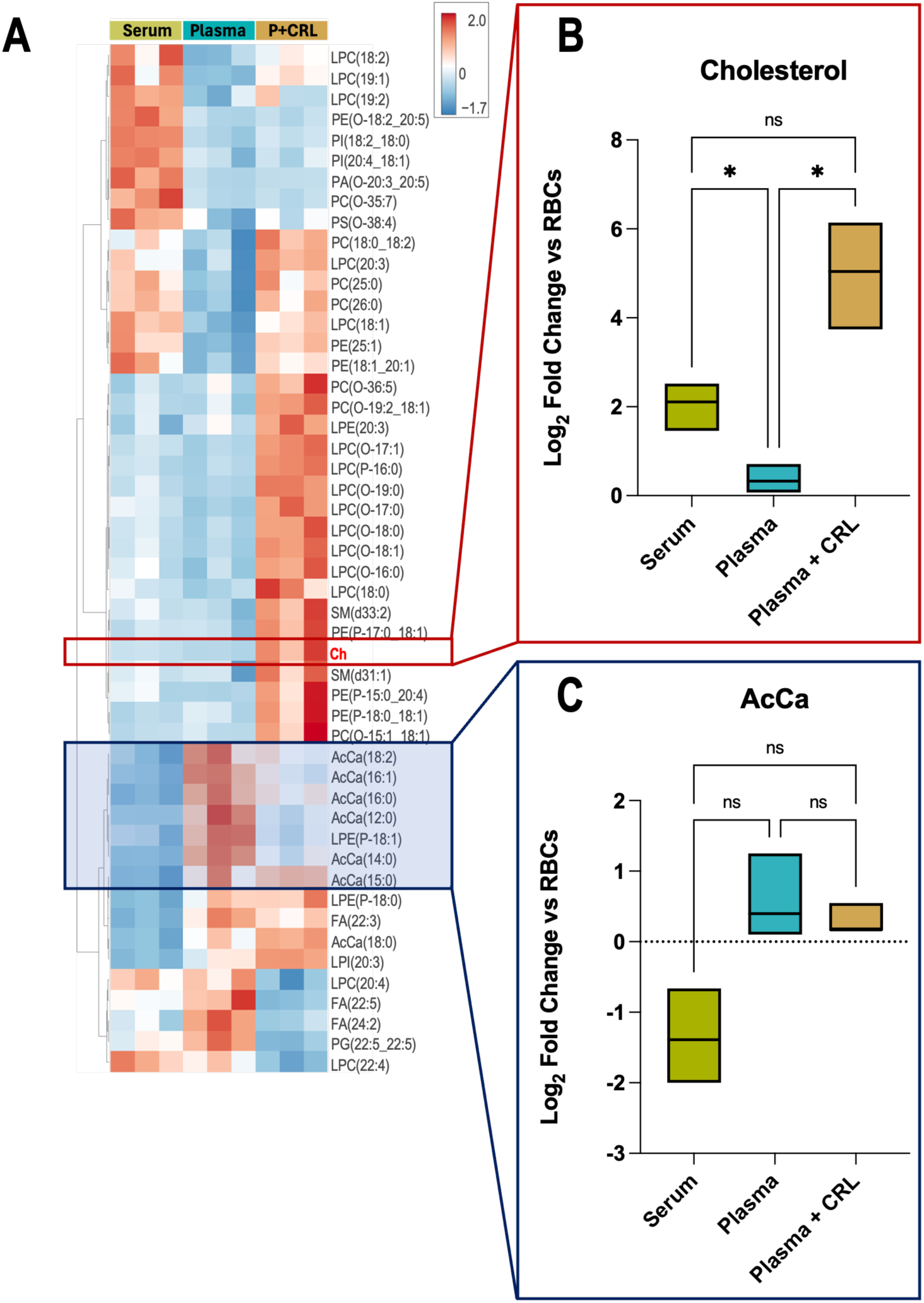
Lipidomic analysis confirms increased cholesterol in serum and supplemented plasma grown primary reticulocytes. **A** Summary heat map of the top 50 significant lipid changes by ANOVA of the lipidome data from serum, plasma, and plasma+CRL cultured reticulocytes in comparison with donor matched erythrocytes. The values represent the average of log_2_(Fold change, FC) of the sum of peak areas of each chain against the equivalent sum in matched-donor red cells. B,C Box plots of Cholesterol (Ch, B) and acyl-carnitines (AcCa, C) log_2_(FC) analysed for significance with non-parametric Kruskal-Wallis test with Dunn’s correction (p<0.05).

### Reduced cholesterol in the plasma-grown reticulocytes impacts reticulocyte characteristics

To assess the properties of the reticulocytes produced using the different lipid sourcesreticulocyte deformability was measured by Automated Rheoscopy (ARCA) ^39^, a single cell ektacytometry system where cells are subjected to a constant and specific shear stress field causing cell elongation. The A/B ratio is used as a deformability measurement, plotted against the frequency of said ratio in the cell population (**Figure 5A**). The weighted average of the A/B ratio was calculated for all 3 donors and the values were plotted in **Figure 5B**. Red blood cells achieve on average a higher A/B ratio, at 1.96 ±0.06 (SD), followed by RET^AB-serum^ with 1.82 ±0.02 (SD), RET^Plasma^ at 1.73 ±0.08 (SD) and RET^Plama+CRL^ at 1.61 ±0.04 (SD). The higher elongation index observed in RBCs compared to cultured reticulocytes is in line with the well-established understanding that mature RBCs are more deformable. Statistically, there is no difference between the means of the three reticulocyte samples but when analysing the distribution peaks in **Figure 4A** there is a visible shift of RET^Plasma+CRL^, indicative of reduced deformability. In fact, comparing the A/B ratio average of cultured reticulocytes with RBCs, only RET^Plasma+CRL^ is substantially reduced, once again indicating a decrease in deformability correlated with the increased cholesterol content.

**Figure 5:**
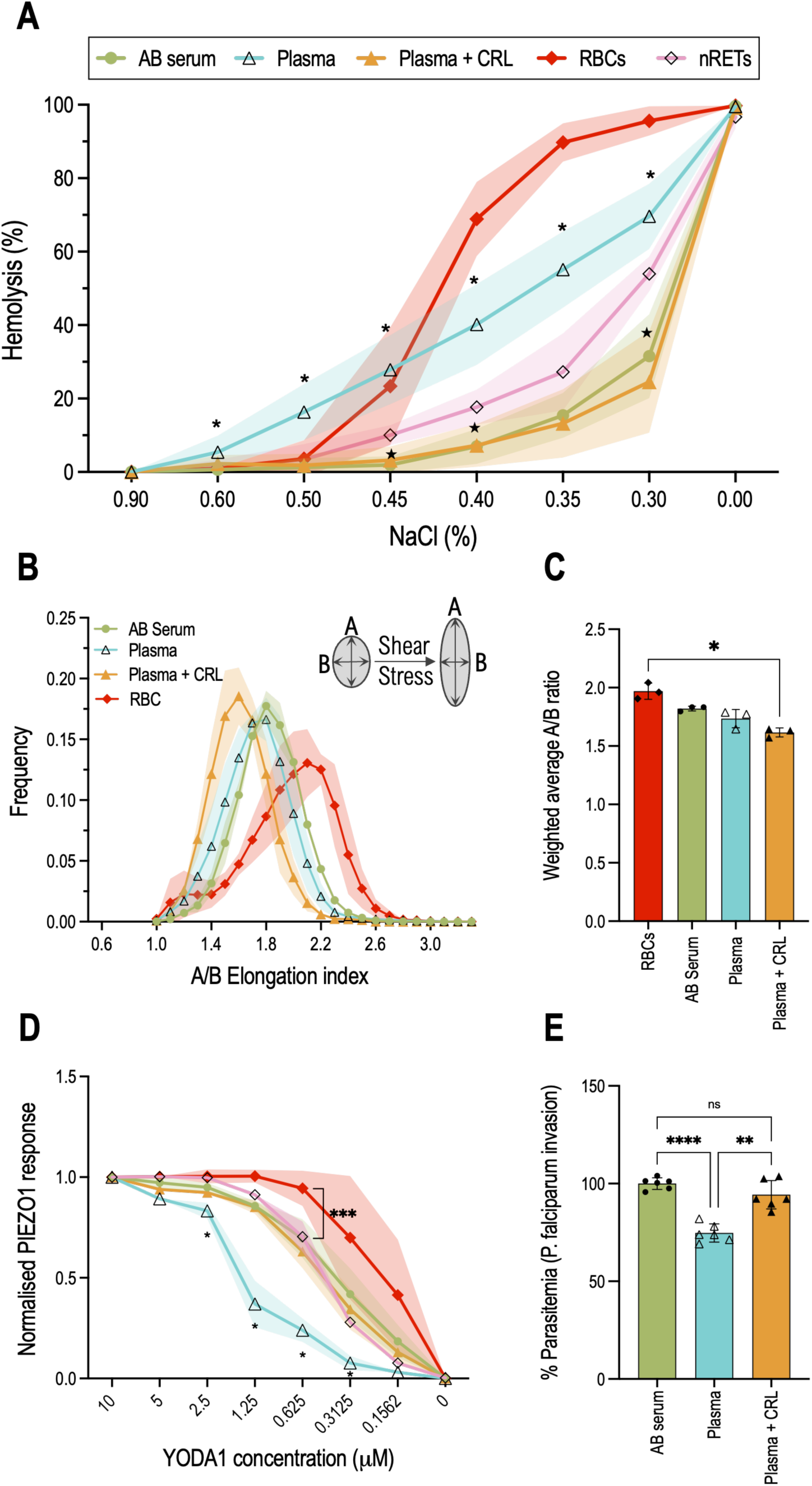
Cholesterol deficiency in cultured reticulocytes modulates deformability, Piezo1 activity and *P. falciparum* invasion. **A** Osmotic fragility analysis of red blood cells (RBCs, n=9), native reticulocytes (nRETs, n=3, CD71+) and filtered reticulocytes cultured in media containing either AB serum (n=6), Plasma (n=6), or Plasma supplemented with Cholesterol-Rich-lipids (n=6). For each donor (n) two technical replicates were performed. Osmotic resistance was calculated based on viable cell counts (flow cytometer) after incubation with decreasing concentrations of NaCl. Non-parametric Mann-Whitney U test with subsequent Dunn correction was performed to test for differences between groups. ∗ p < 0.05 Plasma versus nRETs; ★ p< 0.05 AB serum versus nRETs. **B** Deformability of RBC and cultured reticulocytes measured under shear stress by Automated Rheoscope Cell Analyser, which elongates cells and measures length over width as deformability parameter. (n=3 for each conditions, min of 5000 cells analysed for each). C Weighted average of A/B elongation index for each condition (n=3). Statistical significance was assessed by Kruskal-Wallis test with Dunn’s correction. D Piezo1 activity of RBCs (n=6), nRETs (n=3), and filtered reticulocytes (n=3 each culture condition). Calcium concentration inside the cell is measured pre- and post-YODA1 addition, a chemical inducer of Piezo1 activity, correlating with Piezo1 activity. Data shown is normalised for the highest YODA1 concentration used, 10µM. Multiple t-tests with Welch’s and Holm-Šidak corrections were performed to test for differences between groups. * signifies a p < 0.05 for Plasma versus nRETs and *** equals to p<0.001. E Bar plot quantifying *P. falciparum* reticulocyte invasion efficiency. Invasion was assessed by flow cytometry using a SYBR-green DNA stain (2 separate parasitaemia percentages) and data was normalised to the NT-control (n=3, SD). A parametric Brown-Forsythe and Welch test was performed to test for differences between groups. Plotted data corresponds to the mean with error bars or shaded regions indicating standard deviation. p < 0.05 was considered statistically significant.

As reported previously^24,40,41^, an osmotic fragility test (OFT) can be used to assess reticulocyte fragility. The OFT exposes cells to various dilutions of NaCl, exposing RBCs to a hypotonic environment, causing water to enter the cell, resulting in swelling and ultimately lysis. The OFT was performed on RBCs, native reticulocytes (nRETs) obtained by magnetically isolating CD71+ cells from donor red cells and filtered cultured reticulocytes. As shown in **Figure 5C**, RBCs produce an S-shape curve consistent with 50% haemolysis observed between 0.45% and 0.40% NaCl (w/v) whilst native reticulocytes are more resistant to osmotic changes, requiring a concentration of 0.35% to 0.30% NaCl (w/v) to achieve the same cell lysis percentage. This is consistent with the consensus that reticulocytes are more resistant to osmotic alteration despite being less deformable than RBCs^42^. Both RET^AB-serum^ and RET^Plasma+CRL^ reticulocytes exhibit similar OFT pattern to that observed for nRETs but with significant differences at 0.45%, 0.40% and 0.30% NaCl solutions (w/v). These data indicate a heightened resistance to hypotonic solutions of cultured reticulocytes when compared with nRETs. Plasma-grown reticulocytes display the opposite, an increased fragility even when exposed to high salt concentrations. At 0.5% NaCl (w/v) RET^Plasma^ experience 16.3 ±7.6% haemolysis while only 3.4 ±4.1% of nRETs and 1.2 ±2.4% of RET^AB-serum^ lyse at that same NaCl concentration. This remains a significant difference throughout the remaining NaCl dilutions until all conditions 100% lyse in water.

Multiple reports have highlighted the importance of the lipid environment on the mechanosensory channel Piezo1 properties and activity ^43^ , as such alteration in the different lipid sources and the resulting cholesterol changes could impact Piezo1 function. In order to investigate this RBCs, nRETs, and filtered cultured reticulocytes cultured in the different lipid sources were treated with various concentrations of YODA1. A baseline is first acquired for each sample followed by YODA1 addition which results in a Ca^2+^ influx, detected by Fluo4-AM fluorescence. The YODA1 titration results are plotted in **Figure 5D**, normalised for the highest concentration used (10μM).

Between 10µM and 1.25µM YODA1, no discernible difference in Ca^2+^ influx is observed in erythrocytes, indicating saturating activation of Piezo1. This saturation is visually evident through the plateau in Piezo1 response. The channel activation experiences a gradual decline with decreasing YODA1 concentrations. Comparing nRETs with RBCs there was a difference in response from 1.25μM, with RBCs consistently exhibiting higher Ca^2+^ influxes, also obvious by the required concentration to achieve 50% Piezo1 activation (approximately 0.20μM to 0.46μM). As mentioned before this is to be expected, as reticulocytes are less deformable. Focusing on cultured reticulocytes, there is no difference between nRETs and RET^AB-serum^, indicating that the differences observed between cultured reticulocytes and RBCs are due to their immaturity and not from *ex vivo* erythropoiesis. There is however a detectable decrease of Piezo1 activity observed for RET^Plasma^, that was recovered in full by cholesterol supplementation, as RET^Plasma+CRL^ display the same channel activation levels as nRETs.

Finally, cholesterol also plays a pivotal role in *Plasmodium falciparum* invasion of RBCs as the parasite relies on cholesterol-rich domains in the host cell membrane as entry points. Disruption of cholesterol homeostasis has been shown to impede the invasion process^44^. **Figure 5E** illustrates invasive susceptibility of all three culture conditions (AB-serum, Plasma, Plasma + CRL), with invasions assays performed on filtered reticulocytes and data normalised for RET^AB-serum^. Consistent with the lower cholesterol levels present in RET^Plasma^, a significant reduction in invasion capacity was observed for these reticulocytes. This effect is rescued by cholesterol addition in the RET^Plasma+CRL^ cultured cells.

## Discussion

This study has confirmed that the lipid profile of reticulocytes produced *in vitro* by culturing primary stem cells (CD34^+^) is reassuringly similar to the lipid profiles of native reticulocytes and erythrocytes. It has also confirmed that a significant factor that can influence reticulocyte quality and functionality is the choice of lipid supplementation, particularly the use of human AB-serum versus solvent-treated plasma products such as Octaplas. Our findings show that when using solvent/detergent depleted plasma such as Octaplas, careful optimisation of cholesterol supplementation should be undertaken to ensure that the reticulocytes do not have cholesterol deficiency and so impact on their characteristics. We also show that supplementation of cultures with 3% AB serum provide sufficient cholesterol for erythroid differentiation. Consequently, additional cholesterol supplementation is not required when AB-serum is used. These findings underscore the broader challenge in erythroid culture standardisation, as different laboratories employ varied serum, plasma, or serum-free media formulations, each method will require careful optimisation, characterisation and testing to check for impact on reticulocyte characteristics and functionality.

Despite the challenge of donor biological variability in terms of proliferation, it was noticeable that under the culture conditions using AB-serum as a lipid source may yield better results than plasma alone in terms of proliferation (**Figure 1B**), with cholesterol supplementation further enhancing the proliferation of plasma-grown cells. Given the rapid proliferation of early erythroid progenitors, it is expected that cholesterol availability would impact cell expansion^20,45^. Colabelling with Band 3 and CD49d (**Figure 2C**) corroborated data observed regarding enucleation (**Figure 1C**), where reticulocytes cultured in plasma progressed slower through differentiation. This was independent of cholesterol availability, as CRL supplementation of Plasma cultures did not reverse this effect, marking a key difference between Plasma and Serum. A similar effect was observed regarding CD71 levels, a marker for reticulocyte maturity, with plasma reticulocytes showing reduced expression levels. A curious result given the apparent delayed differentiation observed in the early stages of culture (**Figure 1E**).

Importantly, plasma only cultures exhibit dramatically impaired filtration, producing extremely low recovery yields following leukofiltration (**Figure 1D**). Adding cholesterol-rich lipids to the plasma containing media rescued the reticulocyte filterability to levels observed in AB-serum cultures. This is suggestive of impacted deformability, as successful passage through the leukofilter is highly dependent on cell deformability^32^. A recent study by Claessen et al.^20^ explored the requirement for cholesterol supplementation as an aid to improved filterability and in their study, using plasma as a lipid source. Of note here is the novel correlation between the specific use of plasma in culture media and insufficient cholesterol availability, since erythroid differentiation in the presence of AB-serum does not require cholesterol supplementation. This was corroborated by measuring cholesterol levels in the filtered reticulocytes (**Figure 1F**), which revealed reduced cholesterol in RET^Plasma^ compared with even mature RBCs and is also observed in the comparative lipidomics.

The observed upregulation of *HMGCR*, *SQLE*, and *LDLR* genes in erythroid cells grown in plasma-only conditions is consistent with the hypothesis that the cultured erythroblasts are experiencing cholesterol restriction and so are compensating for the lower cholesterol availability by increasing cellular cholesterol synthesis and uptake. The downregulation of the same genes when cholesterol is supplemented in the culture media is consistent with a feedback mechanism where exogenous cholesterol inhibits the need for intracellular synthesis and uptake. This feedback control is well documented in cholesterol metabolism, where excess cholesterol leads to suppression of biosynthesis and receptor-mediated uptake to maintain homeostasis^46,47^.

SREBP2 mRNA levels also increase in response to low cholesterol conditions, further amplifying the regulatory network that drives the expression of key cholesterol biosynthesis and uptake genes such as HMGCR, SQLE, and LDLR. Interestingly, the observed upregulation of SREBP2 in plasma-only conditions was quite modest, showing less than a two-fold increase, with significant variation between donors. Indeed one donor exhibited no upregulation of the transcription factor and correspondingly low expression of regulated genes HMGCR, SQLE, and LDLR, while the remaining two exhibited more substantial regulation of these genes.

SREBP1 was also significantly upregulated in the same two donors, suggesting that fatty acid metabolism pathways can also be activated alongside cholesterol biosynthesis. In fact, despite SREBP1 being more associated with transcription of fatty acid synthesis genes and SREBP2 more associated with cholesterol-related genes, recent publications have shown how SREBP1 is also activated by low cholesterol and how the upregulation of FASN (Fatty Acid Synthase), a downstream target of SREBP1, has been linked to enhanced cholesterol production^48–51^. Our proteomics analysis of filtered reticulocytes further corroborates this observation, detecting a marked upregulation of FASN in Plasma-cultured cells only. Additionally, the upregulation of FDPS (Farnesyl Diphosphate Synthase) and MVD (Mevalonate Diphosphate Decarboxylase) in the proteomics data indicates an increased flux through the mevalonate pathway, crucial for cholesterol biosynthesis^52^.

Overall, the omics data showed that plasma-grown reticulocytes exhibit a marked activation of lipid and cholesterol biosynthesis pathways, likely driven by SREBP1/2 activity, fuelled by reduced cholesterol depletion. These data also reveal minimal proteomic difference between cultured reticulocytes and mature red blood cells, independent of lipid source used.

We established that RET^AB-serum^ have similar osmotic resistance profile to those of nRETs, whilst RET^Plasma^ showed increased osmotic fragility (**Figure 5A**), consistent with previously published data of cholesterol-deficient reticulocytes levels^24^. The osmotic fragility was reversed by supplementation with CRL. Curiously when submitting the same reticulocytes to an automated rheoscope (**Figure 5B-C**), testing the elongation capacity of the cell, we saw no impairment of the plasma cultured cells, instead the reticulocytes supplemented with extra cholesterol had reduced deformability. Increased membrane stiffness has been described in hypercholesterolemia ^8,53^, aligning with our findings. This suggests that careful optimisation of cholesterol supplementation should be considered, to ensure minimal impact on reticulocyte properties.

Cholesterol not only influences membrane fluidity and stiffness but also regulates the activity of membrane proteins, such as the mechanosensitive ion channel PIEZO1. Studies have shown that PIEZO1 function and activity are dependent on cholesterol-rich lipid rafts^54,55^. By employing a titration of the PIEZO1 agonist YODA1 to cultured reticulocytes (**Figure 5D**), known to reduce the sensitivity threshold of the channel^56^, we observed reduced activity of the channel in cells with reduced cholesterol. These results provide validation of PIEZO1 interaction with cholesterol in human reticulocytes without the necessity of chemically depleting cholesterol, such as with MßCD, a method often used despite producing nonspecific effects and non-physiological cholesterol depletion^57^. In addition to its impact on mechanotransduction, the lipid composition of the membrane is also known to affect interactions with pathogens, with cholesterol playing a key role in the formation of membrane entry points for *Plasmodium falciparum*^58^ .We demonstrated how RET^Plasma^, with low cholesterol content, experience reduced invasion efficiency (**Figure 5E**). Similarly to the other discussed assays, supplementation with cholesterol-rich lipids reversed the effect, rescuing *P. falciparum* invasion to levels observed in serum supplemented cultured reticulocytes, indicating a direct correlation with membrane cholesterol content.

Taken all together, these data underscore the importance of careful selection of lipid supplementation when conducting erythroid culture *ex vivo*. While both AB-serum and plasma can both support erythropoiesis, they do so with distinct impacts on cell maturation, membrane composition, and downstream functionality. These findings have broad implications for the standardisation of *ex vivo* erythropoiesis protocols and the interpretation of functional assays, particularly when reticulocytes are used to study membrane-associated phenomena or host-pathogen interactions.

Future work will need to focus further on dissecting the donor-specific lipid metabolic responses and exploring additional omics-based characterisation to refine serum-free or defined culture systems to ensure that such cultures consistently yield physiologically relevant reticulocytes, especially if these are to be used in humans as a replacement for donor derived RBCs or used to deliver therapeutics.

## Materials and Methods

### Source Material

All human blood material was provided with written informed consent for research use given in accordance with the Declaration of Helsinki (NHSBT, Filton, Bristol). The research into the mechanisms of erythropoiesis was reviewed and approved by Bristol Research Ethics committee (REC Number 12/SW/0199).

### Antibodies, Fluorescent Dyes and Small Molecules

For flow cytometry several antibodies were used. BRIC71, targeting Band 3, is a mouse IgG1 antibody from IBGRL Research Products and was used at a 1:2 dilution. A purified mouse IgG1 isotype control (clone MG1-45) from Biolegend was used at a 1:50 dilution. To detect the transferrin receptor, an APC-conjugated anti-human CD71 antibody (clone CY1G4), a rat IgG2a, was used, at 1:50 dilution. For control, an APC-conjugated anti-mouse IgG1 antibody (clone RMG1-1) was employed, also at 1:50. CD49d was detected using a FITC-conjugated anti-human CD49d antibody (clone MZ18-24A9), a mouse IgG2b from Miltenyi Biotech. A FITC-conjugated mouse IgG2b isotype control (clone IS6-11E5.11) was also included, both at 1:50 dilution.

The following compounds were used for various cellular assays. Filipin III (Sigma-Aldrich, F4767) was applied at 50 μg/mL to stain cholesterol. Fluo-4 (Thermo Fisher Scientific, F14201), a calcium indicator, was used at a concentration of 5 μM. For DNA staining, Hoechst 33342 (Invitrogen, B2261) was used at 5 μg/mL. Thiazole Orange (Sigma-Aldrich, 390062) was employed at 0.1 μg/mL to label RNA. To activate PIEZO1, Yoda1 (Tocris, 5586) was used across a range of concentrations, from 0.156 to 10 μM.

### CD34^+^ culture

Peripheral blood mononuclear cells are isolated from apheresis cones^21^, followed by CD34^+^ magnetic cell isolation with CD34^+^ MicroBead kit (Miltenyi Biotec) according to manufacturer’s protocol. Cells were cultured as described by Kupzig et al ^26^. Isolated cells were seeded at 1 x10^5^/mL in IMDM base medium (3% (v/v) Heat-inactivated (30min at 56°C) Human Male AB Serum (Sigma-Aldrich), 2 mg/mL Human Serum Albumin (HSA; Irvine-Scientific), 10 µg/mL insulin (Sigma-Aldrich), 3 U/mL heparin (Sigma-Aldrich), 0.2 mg/mL holotransferrin (Sigma-Aldrich), 3 U/mL erythropoietin (Epo; Bristol Royal Infirmary), and 100 µg/mL streptomycin (Sigma-Aldrich)). From days 0 to 8 medium was supplemented with 40ng/mL SCF and 1ng/mL IL-3, from days 8-13 only 40ng/mL SCF, and from day 13 onwards no supplementation. Where indicated AB serum and HSA were replaced with 5% (v/v) Octaplas (Octapharma) and/or supplemented with 50mg/L Cholesterol rich lipid mix (L4646 Sigma). Cells were cultured until day 20. To obtain a pure population of reticulocytes, the cultures were filtered by leukofiltration. A leukocyte reduction filter (Macopharma) was pre-soaked and equilibrated with phosphate-buffered saline (PBS) and the cultured cell suspension was loaded into the filter followed by at least three volumes of PBS and allowed to pass through under gravity. The resulting flow-through was then centrifuged at 400 × g, for 15 min and the pelleted cells were resuspended in PBSAG (PBS + 1 mg/ml BSA, 2 mg/ml glucose) and kept at 4 °C if not used immediately.

### Native reticulocyte isolation (CD71+)

CD71+ cells were isolated from the red cell fraction of apheresis waste following density gradient centrifugation. The packed red cell layer was collected, washed 3 times in PBS and magnetic isolation performed using the CD71 MicroBeads cell isolation kit (Miltenyi Biotec) to enrich for CD71-expressing reticulocytes. Briefly, 500μL of packed red cells per donor were washed twice in MACS buffer and resuspended in 5mL buffer. 100μL of anti-CD71 beads were added, and the mixture incubated for 15 minutes at 4°C. After centrifugation, cells were resuspended in 10mL cold MACS buffer and loaded a LS column (maximum 500μL packed red cells per column). At least 7 washes of 5mL MACS buffer were applied and then the cells were eluted in 5mL MACS buffer using the supplied plunger. Purity was assessed by flow cytometry by labelling cells with anti-CD71-APC antibody (Biolegend, San Diego, USA), and the isolated native reticulocytes stored in PBS-AG at 4°C if not used immediately.

### Osmotic Fragility Assays

Filtered reticulocytes (1-2 x10^5^/well) were incubated in decreasing NaCl concentrations (0.9–0%) for 10min at 37°C. Lysis was stopped by adding 4x volume of PBSAG. Live cells, considered as having a normal FSC/SSC profile as defined by the 0.9% NaCl control, were counted by flow cytometry using the MACSQuant10^24^.

### Total cholesterol labelling

Cholesterol labelling^27^ was performed by initially fixing the cells in 1% paraformaldehyde and 0.0075% glutaraldehyde in PBSAG, at room temperature, for 15 minutes, followed by three washed in PBSAG. The fixed cells were incubated with 50μg/mL filipin (from 25mg/mL stock in DMSO, Filipin III from *S. filipinensis*, Sigma-Aldrich) in PBS for 45 minutes in the dark. Cells were washed twice in PBS and analysed on the Fortessa X20 flow cytometer using UV excitation.

### Automated Rheoscopy

A total of 1x10^6^ cells were diluted in 200µL of a polyvinylpyrrolidone solution (viscosity, 28.1mPa·s; Mechatronics Instruments). Cell deformability distributions were assessed in an Automated Rheoscope and Cell Analyzer (ARCA) according to previously published protocols^18^. At least 2000 valid cells per sample were analysed.

### PIEZO1 responsivity assay

Serial dilutions of Yoda-1 (Tocris, Bristol, UK) were prepared from a 20mM stock, and the final assay concentrations used were 10μM, 5μM, 2.5μM, 1.25μM, 625nM, 312.5nM, 156.3nM, and vehicle only (DMSO). 2 x10^5^ of filtered CD34-derived reticulocytes were resuspended in StemSpan containing 5μM of Fluo-4 AM (Thermo Fisher Scientific). Cells were incubated for 45 minutes at 37°C, washed in PBS-AG and resuspended in 100μL IMDM (Source BioScience UK Ltd or Sigma-Aldrich) supplemented with 2% FCS (Gibco). To streamline the assay, for each well 40μL were acquired for a basal fluorescence measurement followed by Yoda-1 addition, using a 100X concentrated stock solution (e.g. adding 0.6μL of 1mM stock to obtain 10μM final concentration), gently mixing the sample, waiting 60 seconds, and measuring the post-PIEZO1 activation calcium influx. The calcium influx was measured using flow cytometry (MACSQuant Analyzer 10)

### Reverse transcription quantitative-PCR (RT-qPCR)

RNA was isolated from frozen pellets (CD34^+^ differentiation day 8) using a RNeasy kit (Qiagen), following supplier’s protocol. Total RNA concentration was measured using NanoDrop (ThermoFisher). Complementary DNA (cDNA) was generated using the QuantiTect Reverse Transcription Kit (Qiagen) following the indicated protocol. The RT-qPCR was done in 20μL reactions in a MicroAmpTM Optical 96-Well Reaction Plate (Applied Biosystems). Assuming a total conversion of RNA to cDNA, 10ng of cDNA was used as template with 10μL of PowerUpTM SYBRTM Green Master Mix (Applied Biosystems), 1μL of each primer (10μM) and 6μL nuclease-free water. Each sample was analysed in triplicate and a no template control was included. The RT-qPCR were performed using the QuantiStudio3^TM^ RT PCR system (Applied Biosystems) under the Standard cycling mode (Primer Tm > 60°C) as indicated by the manufacture’s protocol for SYBR green dyes. The relative gene expression was determined with the 2^-ΛλΛλCt^ method. Primers (Eurofins) used: GAPDH fw GAGTCAACGGATTTGGTCGT; GAPDH rev TTGATTTTGGAGGGATCTCG; HMGCR fw TGGGATGACTCGTGGCCCAGTT; HMGCR rev TGGCATCCCCTGACCTGGACTG; SQLE fw TAAGGAGCAGCTCGAGGCCAGG; SQLE rev CACCCGGCTGCAGGAATTCTCC; LDLR fw CCAACCTGAGGAACGTGGTCGC; LDLR rev AGTGCCCAGGACAGAGTCGGTC; SREBP1 fw TGTGGCGGCTGCATTGAGAGTG; SREBP1 rev GGGGTACTGAGCACGGACCAGT; SREBP2 fw TCGAGTCAGGTTCTGGGGGCTG; SREBP2 rev TGCCTCCAGAAGGTGACCGAGG

### P. falciparum invasion assays

*P. falciparum* strain 3D7 parasites (BEI Resources) were maintained in human erythrocytes at 5% hematocrit using standard culture conditions^1^. Schizont stage parasites were magnetically purified using the Magnetic Cell Separation (MACS) system (Miltenyi Biotec) and added to wells of a round bottomed 96-well plate containing 1x10^6^ leukofiltered CD34-derived reticulocytes. Parasitemias ranging from 1-8% were used. Heparin (100 mU/µl final) was used to inhibit invasion in negative controls. After ∼18 h, invasion was quantified using flow cytometry as previously described^2,3^. For flow cytometry, cells were stained with SYBR Green (1:2000 in culture media; Sigma-Aldrich) for 30 min at 37°C in the dark. Cells were centrifuged, SYBR Green containing media removed and then fixed for 15 minutes at room temperature as described above. Invasion was quantified based on SYBR green positivity using the heparin control to correct for background events within the invasion gate.

### Metabolomics

Metabolomics analyses were performed as previously described^4^. Cells were extracted in ice cold 5:3:2 MeOH:MeCN:water (v/v/v) at a 1x10^6^ cells/ml ratio, then vortexed for 30 min at 4 °C. Supernatants were clarified by centrifugation (10 min, 12,000 g, 4 °C). The resulting metabolite extracts were analysed (10 uL per injection) by ultra-high-pressure liquid chromatography coupled to mass spectrometry (UHPLC-MS — Vanquish and QExactive, Thermo). Metabolites were resolved on a Phenomenex Kinetex C18 column (2.1 x 150 mm, 1.7 um) at 45 °C using a 5-minute gradient method in positive and negative ion modes (separate runs) over the scan range 65-975 m/z exactly as previously described^4^. Oxylipins were resolved on a Waters ACQUITY UPLC BEH C18 column (2.1 x 100 mm, 1.7 µm) at 60 °C using mobile phase (A) of 20:80:0.02 MeCN:water:formic acid (FA) and a mobile phase (B) of 20:80:0.02 MeCN:isopropanol:FA. For negative mode analysis the chromatographic the gradient was as follows: 0.35 mL/min flowrate, 0% B 0-0.5 min, 25% B at 1 min, 40% B at 2.5min, 55% B at 2.6min, 70% B at 4.5 min, 100% B at 4.6-6 min, 0% B at 6.1-7 min. The Q Exactive MS was operated in negative ion mode, scanning in Full MS mode (2 μscans) from 150 to 1500 m/z at 70,000 resolution, with 4 kV spray voltage, 45 sheath gas, 15 auxiliary gas. Following data acquisition, .raw files were converted to .mzXML using RawConverter then metabolites assigned and peaks integrated using Maven (Princeton University) in conjunction with the KEGG database and an in-house standard library. Quality control was assessed as using technical replicates run at beginning, end, and middle of each sequence as previously described^5,6^.

### Lipidomics

Total lipids were extracted as previously described^7^: cells were extracted in cold methanol at a 1x10^6^ cells/ml ratio. Samples were then briefly vortexed and incubated at -20 °C for 30 minutes. Following incubation, samples were centrifuged at 12,700 RPM for 10 minutes at 4 °C and 80 μL of supernatant was transferred to a new tube for analysis. Lipid extracts were analyzed (10 uL per injection) on a Thermo Vanquish UHPLC/Q Exactive MS system using a 5 min lipidomics gradient and a Kinetex C18 column (30 x 2.1 mm, 1.7 µm, Phenomenex) held at 50 °C. Mobile phase A: 25:75 MeCN:water with 5 mM ammonium acetate; Mobile phase B: 90:10 isopropanol:MeCN with 5 mM ammonium acetate. The gradient and flow rate were as follows: 0.3 mL/min of 10% B at 0 min, 0.3 mL/min of 95% B at 3 min, 0.3 mL/min of 95% B at 4.2 min, 0.45 mL/min 10% B at 4.3 min, 0.4 mL/min of 10% B at 4.9 min, and 0.3 mL/min of 10% B at 5 min. Samples were run in positive and negative ion modes (both ESI, separate runs) at 125 to 1500 m/z and 70,000 resolution, 4 kV spray voltage, 45 sheath gas, 25 auxiliary gas. The MS was run in data-dependent acquisition mode (ddMS^2^) with top10 fragmentation. Raw MS data files were searched using LipidSearch v 5.0 (Thermo).

### Proteomics

Proteomics analyses were performed as described^8^. A volume of 10 μL of cells were lysed in 90 μL of distilled water; 5 μL of lysates were mixed with 45 μL of 5% SDS and then vortexed. Samples were reduced with 10 mM DTT at 55 °C for 30 min, cooled to room temperature, and then alkylated with 25 mM iodoacetamide in the dark for 30 min. Next, a final concentration of 1.2% phosphoric acid and then six volumes of binding buffer (90% methanol; 100 mM triethylammonium bicarbonate, TEAB; pH 7.1) were added to each sample. After gentle mixing, the protein solution was loaded to a S-Trap 96-well plate, spun at 1500 x g for 2 min, and the flow-through collected and reloaded onto the 96-well plate. This step was repeated three times, and then the 96-well plate was washed with 200 μL of binding buffer 3 times. Finally, 1 μg of sequencing-grade trypsin (Promega) and 125 μL of digestion buffer (50 mM TEAB) were added onto the filter and digested carried out at 37 °C for 6 h. To elute peptides, three stepwise buffers were applied, with 100 μL of each with one more repeat, including 50 mM TEAB, 0.2% formic acid (FA), and 50% acetonitrile and 0.2% FA. The peptide solutions were pooled, lyophilized, and resuspended in 500 μL of 0.1 % FA.

Each sample was loaded onto individual Evotips for desalting and then washed with 200 μL 0.1% FA followed by the addition of 100 μL storage solvent (0.1% FA) to keep the Evotips wet until analysis. The Evosep One system (Evosep, Odense, Denmark) was used to separate peptides on a Pepsep column, (150 um inter diameter, 15 cm) packed with ReproSil C18 1.9 um, 120A resin. The system was coupled to a timsTOF Pro mass spectrometer (Bruker Daltonics, Bremen, Germany) via a nano-electrospray ion source (Captive Spray, Bruker Daltonics). The mass spectrometer was operated in PASEF mode. The ramp time was set to 100 ms and 10 PASEF MS/MS scans per topN acquisition cycle were acquired. MS and MS/MS spectra were recorded from *m/z* 100 to 1700. The ion mobility was scanned from 0.7 to 1.50 Vs/cm2. Precursors for data-dependent acquisition were isolated within ± 1 Th and fragmented with an ion mobility-dependent collision energy, which was linearly increased from 20 to 59 eV in positive mode. Low-abundance precursor ions with an intensity above a threshold of 500 counts but below a target value of 20000 counts were repeatedly scheduled and otherwise dynamically excluded for 0.4 min.

### Database Searching and Protein Identi>cation

MS/MS spectra were extracted from raw data files and converted into .mgf files using MS Convert (ProteoWizard, v. 3.0). Peptide spectral matching was performed with Mascot (v. 2.5) against the Uniprot human database. Mass tolerances were +/-15 ppm for parent ions, and +/-0.4 Da for fragment ions. Trypsin specificity was used, allowing for 1 missed cleavage. Protein N-terminal acetylation, isopeptide bond formation with loss of ammonia (K), and peptide N-terminal pyroglutamic acid formation were set as variable modifications with Cys carbamidomethylation set as a fixed modification.

Scaffold (v 4.8, Proteome Software, Portland, OR, USA) was used to validate MS/MS based peptide and protein identifications. Peptide identifications were accepted if they could be established at greater than 95.0% probability as specified by the Peptide Prophet algorithm. Protein identifications were accepted if they could be established at greater than 99.0% probability and contained at least two identified unique peptides.

### Statistics

Data was organised and analysed using either Excel or GraphPad Prism 10 software. Unless stated otherwise, data is displayed as mean ± standard deviation (SD). Statistical analysis was completed where appropriate, first verifying normal distribution of the data using the Shapiro-Wilk normality test (significance level of 0.05). For non-parametric data sets, the Mann-Whitney U test was used to compare 2 groups and a Kruskal-Wallis test with Bonferroni correction was used when comparing 3 or more groups. For parametric data sets a Brown-Forsythe and Welch test was performed to test for differences between groups. P < 0.05 was set as statistical significance threshold (*p < 0.05, **p < 0.01, ***p < 0.001. ****p < 0.0001.

For omics data, statistical analysis were conducted using MetaboAnalyst v 5.0 and Rstudio upon autoscale normalization (i.e., data were mean-centred, divided by the standard deviation of each variable). Line plots and volcano plots of correlations (Spearman) were generated upon analysis of the raw data via Rstudio. Significance was calculated upon false discovery rate correction. Network analyses and pathway analyses were performed in OmicsNet v 2.0 using as input the significantly altered metabolites and proteins (FDR-corrected p < 0.05).

## Supporting information

Freire et al 2025 Supplemental figure 1

## Acknowledgements

This study was supported through funding provided by the European Union ITN ‘EVIDENCE’ grant agreement ID 860436 for CMF, the Medical Research Council (MR/V010506/1) for TJS, NRK and infrastructure support funding from the National Institute for Health Research Blood and Transplant Research Unit (NIHR BTRU) in Red Cell Products (IS-BTU-1214-10032). AD was supported by funds from the National Heart, Lung and Blood Institutes (NHLBI) R01 HL146442, R01 HL161004, R01 HL148151, R21 HL150032

The views expressed are those of the authors and not necessarily those of the National Health Service, NIHR, or the Department of Health and Social Care

## Author contributions

CMF, AMT and TJS conceived and designed the study. CMF performed and analysed most experiments, NRK contributed to experimental work, including the planning and execution of the *P. falciparum* assay. JGGD and GJS developed ARCA hardware and analysis software. MD, DS, and AD conducted omics analyses and interpretation. AMT and TJS supervised the study. CMF and AMT wrote the manuscript. All authors reviewed and approved the final manuscript.

## Competing Interests Statement

AMT is a co-founder, a Director and consultant to Scarlet Therapeutics Ltd. TJS is a co-founder and scientific consultant to Scarlet Therapeutics Ltd.

## Data availability

The proteomics data set is available at MassIVE. The MassIVE identifier is MSV000094204, and can be accessed directly through the link https://massive.ucsd.edu/ProteoSAFe/private-dataset.jsp?task=ab88082fec2b43b2952a16535b013127; the metabolomics and lipidomics data sets are available at the National Institutes of Health Common Fund’s National Metabolomics Data Repository website, the Metabolomics Workbench, https://www.metabolomicsworkbench.org for which it has been assigned study ID ST003108.

## Notes

### Competing Interest Statement

The authors have declared no competing interest.

## References

1. Moreau A, Yaya F, Lu H, et al. Physical mechanisms of red blood cell splenic filtration. Proc Natl Acad Sci U S A. 2023;120(44):e2300095120. doi:10.1073/PNAS.2300095120/SUPPL_FILE/PNAS.2300095120.SM06.AVI

2. Minetti G, Dorn I, Köfeler H, Perotti C, Kaestner L. Insights from lipidomics into the terminal maturation of circulating human reticulocytes. Cell Death Discovery 2025 11:1. 2025;11(1):1-12. doi:10.1038/s41420-025-02318-x

3. Mohandas N, Evans E. Mechanical properties of the red cell membrane in relation to molecular structure and genetic defects. Annu Rev Biophys Biomol Struct. 1994;23(Volume 23, 1994):787–818. doi:10.1146/ANNUREV.BB.23.060194.004035/CITE/REFWORKS

4. RA C. Abnormalities of cell-membrane fluidity in the pathogenesis of disease. N Engl J Med. 1977;297:371–377. Accessed September 17, 2024. https://cir.nii.ac.jp/crid/1570572700582030592

5. Paukner K, Lesná IK, Poledne R. Cholesterol in the Cell Membrane—An Emerging Player in Atherogenesis. International Journal of Molecular Sciences 2022, Vol 23, Page 533. 2022;23(1):533. doi:10.3390/IJMS23010533

6. Buchwald H, O’Dea TJ, Menchaca HJ, Michalek VN, Rohde TD. Effect Of Plasma Cholesterol On Red Blood Cell Oxygen Transport. Clin Exp Pharmacol Physiol. 2000;27(12):951–955. doi:10.1046/J.1440-1681.2000.03383.X

7. Forsyth AM, Braunmüller S, Wan J, Franke T, Stone HA. The effects of membrane cholesterol and simvastatin on red blood cell deformability and ATP release. Microvasc Res. 2012;83(3):347–351. doi:10.1016/J.MVR.2012.02.004

8. Doole FT, Kumarage T, Ashkar R, Brown MF. Cholesterol Stiffening of Lipid Membranes. The Journal of Membrane Biology 2022 255:4. 2022;255(4):385-405. doi:10.1007/S00232-022-00263-9

9. Nemkov T, Kingsley PD, Dzieciatkowska M, et al. Circulating primitive murine erythroblasts undergo complex proteomic and metabolomic changes during terminal maturation. Blood Adv. 2022;6(10):3072–3089. doi:10.1182/BLOODADVANCES.2021005975

10. Moon SH, Huang CH, Houlihan SL, et al. p53 Represses the Mevalonate Pathway to Mediate Tumor Suppression. Cell. 2018;176(3):564. doi:10.1016/J.CELL.2018.11.011

11. D’Alessandro A, Keele GR, Hay A, et al. Ferroptosis regulates hemolysis in stored murine and human red blood cells. Blood. 2025;145(7):765–783. doi:10.1182/BLOOD.2024026109

12. Mahdi A, Wodaje T, Kövamees O, et al. The red blood cell as a mediator of endothelial dysfunction in patients with familial hypercholesterolemia and dyslipidemia. J Intern Med. 2022;293(2):228. doi:10.1111/JOIM.13580

13. Himbert S, Qadri SM, Sheffield WP, Schubert P, D’Alessandro A, Rheinstädter MC. Blood bank storage of red blood cells increases RBC cytoplasmic membrane order and bending rigidity. PLoS One. 2021;16(11):e0259267. doi:10.1371/JOURNAL.PONE.0259267

14. Peltier S, Marin M, Dzieciatkowska M, et al. Proteostasis and metabolic dysfunction characterize a subset of storage-induced senescent erythrocytes targeted for post-transfusion clearance. J Clin Invest. Published online March 11, 2025. doi:10.1172/JCI183099

15. Hawksworth J, Satchwell TJ, Meinders M, et al. Enhancement of red blood cell transfusion compatibility using CRISPR-mediated erythroblast gene editing. EMBO Mol Med. 2018;10(6):e8454. doi:10.15252/emmm.201708454

16. Giarratana MC, Rouard H, Dumont A, et al. Proof of principle for transfusion of in vitro-generated red blood cells. Blood. 2011;118(19):5071–5079. doi:10.1182/BLOOD-2011-06-362038

17. Wilson MC, Trakarnsanga K, Heesom KJ, et al. Comparison of the proteome of adult and cord erythroid cells, and changes in the proteome following reticulocyte maturation. Molecular and Cellular Proteomics. 2016;15(6):1938–1946. doi:10.1074/mcp.M115.057315

18. Moura PL, Hawley BR, Mankelow TJ, et al. Non-muscle myosin II drives vesicle loss during human reticulocyte maturation. Haematologica. 2018;103(12):1997–2007. doi:10.3324/HAEMATOL.2018.199083

19. Bernecker C, Lima M, Kolesnik T, et al. Biomechanical properties of native and cultured red blood cells–Interplay of shape, structure and biomechanics. Front Physiol. 2022;13:979298. doi:10.3389/FPHYS.2022.979298/FULL

20. Claessen MJAG, Yagci N, Fu K, et al. Production and stability of cultured red blood cells depends on the concentration of cholesterol in culture medium. Scientific Reports 2024 14:1. 2024;14(1):1-13. doi:10.1038/s41598-024-66440-z

21. Satchwell TJ, Wright KE, Haydn-Smith KL, et al. Genetic manipulation of cell line derived reticulocytes enables dissection of host malaria invasion requirements. Nat Commun. 2019;10(1). doi:10.1038/S41467-019-11790-W

22. Martins Freire C, King NR, Dzieciatkowska M, et al. Complete absence of GLUT1 does not impair human terminal erythroid differentiation. Blood Adv. 2024;8(19):5166–5178. doi:10.1182/BLOODADVANCES.2024012743/2232404/BLOODADVANCES.2024012743.PDF

23. Moura PL, Hawley BR, Dobbe JGG, et al. PIEZO1 gain-of-function mutations delay reticulocyte maturation in hereditary xerocytosis. Haematologica. 2020;105(6):e268. doi:10.3324/HAEMATOL.2019.231159

24. Bernecker C, Köfeler H, Pabst G, et al. Cholesterol Deficiency Causes Impaired Osmotic Stability of Cultured Red Blood Cells. Front Physiol. 2019;10:1529. doi:10.3389/fphys.2019.01529

25. Zingariello M, Bardelli C, Sancillo L, et al. Dexamethasone Predisposes Human Erythroblasts Toward Impaired Lipid Metabolism and Renders Their ex vivo Expansion Highly Dependent on Plasma Lipoproteins. Front Physiol. 2019;10(APR):281. doi:10.3389/fphys.2019.00281

26. Kupzig S, Parsons SF, Curnow E, Anstee DJ, Blair A. Superior survival of ex vivo cultured human reticulocytes following transfusion into mice. Haematologica. 2017;102(3):476–483. doi:10.3324/haematol.2016.154443

27. Maxfield FR, Wüstner D. Analysis of cholesterol trafficking with fluorescent probes. Methods Cell Biol. 2012;108:367. doi:10.1016/B978-0-12-386487-1.00017-1

28. Thomas T, Stefanoni D, Dzieciatkowska M, et al. Evidence for structural protein damage and membrane lipid remodeling in red blood cells from COVID-19 patients. J Proteome Res. 2020;19(11):4455. doi:10.1021/ACS.JPROTEOME.0C00606

29. Nemkov T, Reisz JA, Gehrke S, Hansen KC, D’Alessandro A. High-throughput metabolomics: Isocratic and gradient mass spectrometry-based methods. Methods in Molecular Biology. 2019;1978:13–26. doi:10.1007/978-1-4939-9236-2_2/TABLES/1

30. Reisz JA, Zheng C, D’Alessandro A, Nemkov T. Untargeted and semi-targeted lipid analysis of biological samples using mass spectrometry-based metabolomics. Methods in Molecular Biology. 2019;1978:121–135. doi:10.1007/978-1-4939-9236-2_8/FIGURES/1

31. Griffiths RE, Kupzig S, Cogan N, et al. Maturing reticulocytes internalize plasma membrane in glycophorin A-containing vesicles that fuse with autophagosomes before exocytosis. Published online 2012. doi:10.1182/blood-2011-09-376475

32. Bruil A, Beugeling T, Feijen J, van Aken WG. The mechanisms of leukocyte removal by filtration. Transfus Med Rev. 1995;9(2):145–166. doi:10.1016/S0887-7963(05)80053-7

33. Malleret B, Xu F, Mohandas N, et al. Significant Biochemical, Biophysical and Metabolic Diversity in Circulating Human Cord Blood Reticulocytes. PLoS One. 2013;8(10):e76062. doi:10.1371/journal.pone.0076062

34. Hu J, Liu J, Xue F, et al. Isolation and functional characterization of human erythroblasts at distinct stages: implications for understanding of normal and disordered erythropoiesis in vivo. Blood. 2013;121(16):3246–3253. doi:10.1182/blood-2013-01-476390

35. Gautier EF, Leduc M, Cochet S, et al. Absolute proteome quantification of highly purified populations of circulating reticulocytes and mature erythrocytes. Published online 2018. doi:10.1182/bloodadvances.2018023515

36. Anderson ME, Meister A. Transport and direct utilization of gamma-glutamylcyst(e)ine for glutathione synthesis. Proc Natl Acad Sci U S A. 1983;80(3):707. doi:10.1073/PNAS.80.3.707

37. Maiorino FM, Brigelius-Flohé R, Aumann KD, Roveri A, Schomburg D, Flohé L. Diversity of glutathione peroxidases. Methods Enzymol. 1995;252(C):38–48. doi:10.1016/0076-6879(95)52007-4

38. Stevens-Hernandez CJ, Flatt JF, Kupzig S, Bruce LJ. Reticulocyte Maturation and Variant Red Blood Cells. Front Physiol. 2022;13. doi:10.3389/FPHYS.2022.834463/FULL

39. Kalfa TA. Diagnosis and clinical management of red cell membrane disorders. Hematology. 2021;2021(1):331–340. doi:10.1182/HEMATOLOGY.2021000265

40. Domingues CC, Ciana A, Buttafava A, et al. Effect of cholesterol depletion and temperature on the isolation of detergent-resistant membranes from human erythrocytes. Journal of Membrane Biology. 2010;234(3):195–205. doi:10.1007/S00232-010-9246-5/FIGURES/4

41. Sagawa S, Shirakii K. CHANGES OF OSMOTIC FRAGILITY OF RED BLOOD CELLS DUE TO REPLETION OR DEPLETION OF CHOLESTEROL IN HUMAN AND RAT RED CELLS IN VITRO. J Nutr Sci Vitaminol. 1980;26:161–169.

42. Nagasawa T. Deformability and osmotic fragility of phenylhydrazine-injected rat erythrocytes fractionated by Percoll density-gradients. Jpn J Physiol. 1982;32(2):161–170. doi:10.2170/JJPHYSIOL.32.161

43. Vasileva V, Chubinskiy-Nadezhdin V. Regulation of PIEZO1 channels by lipids and the structural components of extracellular matrix/cell cytoskeleton. J Cell Physiol. 2023;238(5):918–930. doi:10.1002/JCP.31001

44. Geoghegan ND, Evelyn C, Whitehead LW, et al. 4D analysis of malaria parasite invasion offers insights into erythrocyte membrane remodeling and parasitophorous vacuole formation. Nature Communications 2021 12:1. 2021;12(1):1-16. doi:10.1038/s41467-021-23626-7

45. Lu Z, Huang L, Li Y, et al. Fine-Tuning of Cholesterol Homeostasis Controls Erythroid Differentiation. Published online 2021. doi:10.1002/advs.202102669

46. Luo J, Yang H, Song BL. Mechanisms and regulation of cholesterol homeostasis. doi:10.1038/s41580-019-0190-7

47. Duan Y, Gong K, Xu S, Zhang F, Meng X, Han J. Regulation of cholesterol homeostasis in health and diseases: from mechanisms to targeted therapeutics. Signal Transduction and Targeted Therapy 2022 7:1. 2022;7(1):1-29. doi:10.1038/s41392-022-01125-5

48. Du Q, Liu P, Zhang C, et al. FASN promotes lymph node metastasis in cervical cancer via cholesterol reprogramming and lymphangiogenesis. Cell Death & Disease 2022 13:5. 2022;13(5):1-14. doi:10.1038/s41419-022-04926-2

49. Jin Y, Chen Z, Dong J, et al. SREBP1/FASN/cholesterol axis facilitates radioresistance in colorectal cancer. FEBS Open Bio. 2021;11(5):1343–1352. doi:10.1002/2211-5463.13137

50. Carroll RG, Zasłona Z, Galván-Peña S, et al. An unexpected link between fatty acid synthase and cholesterol synthesis in proinflammatory macrophage activation. Journal of Biological Chemistry. 2018;293(15):5509–5521. doi:10.1074/jbc.RA118.001921

51. Geng F, Zhong Y, Su H, et al. SREBP-1 upregulates lipophagy to maintain cholesterol homeostasis in brain tumor cells. Cell Rep. 2023;42(7):112790. doi:10.1016/J.CELREP.2023.112790/ATTACHMENT/7DD0985A-F539-4841-8AC1-2B1E08EA0EEF/MMC9.PDF

52. Brown AJ, Coates HW, Sharpe LJ. Cholesterol synthesis. Biochemistry of Lipids, Lipoproteins and Membranes. Published online January 1, 2021:317–355. doi:10.1016/B978-0-12-824048-9.00005-5

53. Koter M, Franiak I, Strychalska K, Broncel M, Chojnowska-Jezierska J. Damage to the structure of erythrocyte plasma membranes in patients with type-2 hypercholesterolemia. Int J Biochem Cell Biol. 2004;36(2):205–215. doi:10.1016/S1357-2725(03)00195-X

54. Buyan A, Allender DW, Corry B, Schick M. Lipid redistribution in the highly curved footprint of Piezo1. Biophys J. 2023;122(11):1900–1913. doi:10.1016/j.bpj.2022.07.022

55. Beverley KM, Levitan I. Cholesterol regulation of mechanosensitive ion channels. Front Cell Dev Biol. 2024;12:1352259. doi:10.3389/FCELL.2024.1352259/BIBTEX

56. Botello-Smith WM, Jiang W, Zhang H, et al. A mechanism for the activation of the mechanosensitive Piezo1 channel by the small molecule Yoda1. Nature Communications 2019 10:1. 2019;10(1):1-10. doi:10.1038/s41467-019-12501-1

57. Ridone P, Pandzic E, Vassalli M, et al. Disruption of membrane cholesterol organization impairs the activity of PIEZO1 channel clusters. Journal of General Physiology. 2020;152(8). doi:10.1085/JGP.201912515/VIDEO-4

58. Samuel BU, Mohandas N, Harrison T, et al. The Role of Cholesterol and Glycosylphosphatidylinositol-anchored Proteins of Erythrocyte Rafts in Regulating Raft Protein Content and Malarial Infection. Journal of Biological Chemistry. 2001;276(31):29319–29329. doi:10.1074/jbc.M101268200

