## Supplementary material for "Choice of lipid supplementation for *in vitro* erythroid cell culture impacts reticulocyte yield and characteristics": Freire et al 2025 Supplemental figure 1

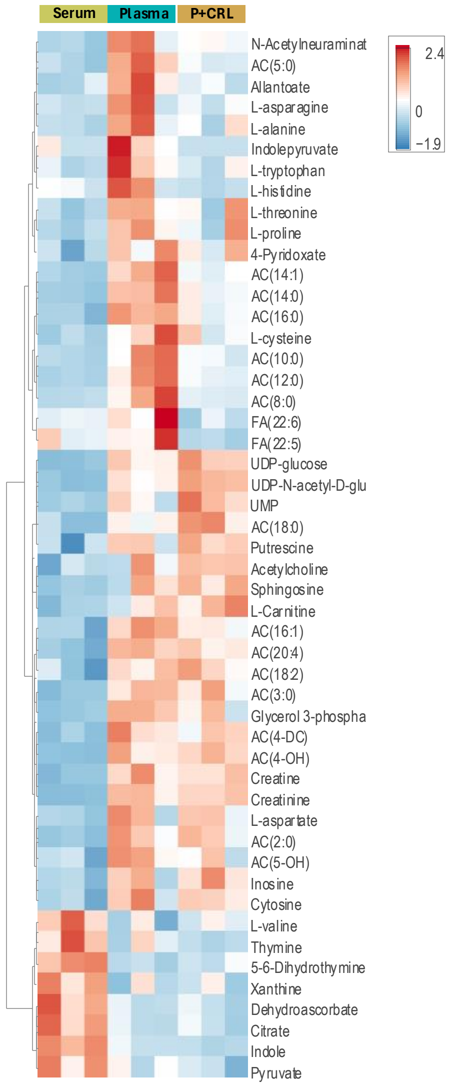


**Supplemental figure 1: Summary of metabolic differences in CD34+ cell-derived reticulocytes grown in the presence of Serum, Plasma, or Plasma supplemented with cholesterol-rich lipids**

Serum, Plasma and Plasma+CRL compared with control RBCs (top 50 metabolites by ANOVA).
